# X-SPATIO: An Explanatory Deep Learning Pipeline for the Prediction and Visualization of Spatially Resolved Biomarker Expression in Triple-Negative Breast Cancer

**DOI:** 10.64898/2026.02.09.704587

**Authors:** Vibha R. Rao, Madhumala K. Sadanandappa, Candice C. Black, Scott M. Palisoul, Adrienne A. Workman, Todd A. MacKenzie, Xiaoying Liu, Mary D. Chamberlin, Louis J. Vaickus, George J. Zanazzi, Shrey S. Sukhadia

## Abstract

Histopathologic evaluation remains central to cancer diagnosis and treatment planning, yet the molecular programs underlying distinct tissue morphologies are not routinely accessible in clinical workflows. Spatial transcriptomic/proteomic platforms provide region-specific molecular measurements but are limited by cost, throughput, and scalability. Most computational pathology models rely on either bulk tissue-based gene expression or a focused gene/protein expression-panel prediction, thereby obscuring subregion-specific morpho–molecular relationships and limiting spatial interpretation of a wider gene/protein expression network. This limitation is particularly significant in triple-negative breast cancer (TNBC), which exhibits pronounced spatial heterogeneity across tumor, stroma, and immune compartments. We developed X-SPATIO, a spatially compatible computational pipeline designed to directly link hematoxylin and eosin (H&E) morphology with region-matched mRNA and protein expression. The model was trained on H&E-defined regions of interest paired with spatially-resolved transcriptomic and proteomic data obtained from GeoMx Digital Spatial Profiler. Using a multiple-instance learning approach, X-SPATIO captures morpho–molecular associations, generating spatio-morphologic attention maps that indicate predictive tissue regions. X-SPATIO demonstrated strong performance across biologically relevant spatial biomarkers, achieving area under the curve values ranging [0.79, 0.97]. Attention maps revealed spatial patterns consistent with known biology, indicating alignment between learned features and tissue organization. By integrating spatial molecular ground truth with routine histopathology, X-SPATIO enables cost-effective inference of spatial biomarker expression and establishes a foundation for biologically grounded discovery and precision oncology in TNBC.

## INTRODUCTION

Histopathologic evaluation remains the foundation of cancer diagnosis, grading, and therapeutic decision-making, yet the molecular programs that give rise to distinct tissue morphologies are not routinely accessible in clinical practice ^1^. Recent advances in spatial transcriptomics and proteomics, such as GeoMx Digital Spatial Profiling (DSP), CosMx,Visium, Xenium, MERFISH, seqFISH, and others, now enable spatially-resolved measurement of gene/protein (biomarker) expression that reflects these underlying molecular programs while preserving tissue context within the tumor and its microenvironment ^2-4^. Although these platforms provide substantial biological insight, their high cost, specialized instrumentation, and operational complexity continue to limit routine clinical adoption and large-scale research deployment ^5^.

In parallel, computational pathology has demonstrated that standard hematoxylin and eosin (H&E)-stained tissue slides contain abundant, quantifiable morphological signals predictive of transcriptional activity, protein expression patterns, microenvironmental organization, and clinically actionable phenotypes ^6,7^. Deep learning and multiple-instance learning (MIL) models provide the primary framework for learning these associations, achieving strong performance across tasks ^8-10^, while attention-based MIL approaches further introduce spatial interpretability by localizing regions that drive model predictions ^11,12^. However, most existing models rely on either bulk tissue-based gene expression or a focused gene/protein expression-panel prediction, which not only limits the ability to determine how well the learned morphological patterns correspond to underlying spatial biology but also limits spatial interpretation of a wider gene/protein expression network ^13,14^. Without spatially-resolved molecular measurements to anchor supervision, these approaches lack direct region-level validation of predicted molecular signals, constraining biological interpretability and mechanistic insight. While platforms such as 10x Genomics Visium and Xenium offer high-resolution spatial measurements, their cost and limited tissue throughput per slide can restrict large-scale modeling efforts. In contrast, NanoString (now Bruker) GeoMx DSP enables profiling of a substantially larger number of tissues within a single slide, supporting rapid and cost-effective generation of spatial data for computational modeling ^15^. In addition, GeoMx DSP supports multiomic spatial analysis, enabling simultaneous profiling of RNA and protein expression, and offers the highest-plex commercially available antibody-based spatial proteomics panels, allowing comprehensive characterization of tissue biology at scale ^15^.

These advances become significant particularly for triple-negative breast cancer (TNBC), a generally aggressive breast cancer subtype lacking expression of estrogen receptor, progesterone receptor, and human epidermal growth factor receptor 2 (HER2), where limited targeted therapies underscore the need for additional biomarkers to guide treatment and prognosis. TNBC is characterized by profound intratumoral heterogeneity, rapid evolution, and complex interactions among tumor, stroma, and immune components ^16,17^. Biomarker expression, such as those governing epithelial-mesenchymal transition (EMT), stromal remodeling, immune activation, and cell-cycle dysregulation, strongly influence TNBC prognosis and therapeutic response, yet these biomarkers manifest in spatially distinct morphologic niches^16^. Consequently, tools that can spatially connect H&E-derived morphology with molecular phenotype are required to significantly deepen biological understanding and guide biomarker discovery in TNBC.

To address this gap, we present X-SPATIO, a focused pipeline for learning and localizing molecular signals from tissue morphology using spatial proteogenomic data. X-SPATIO is trained using H&E and morphology marker-defined regions of interest (ROIs), where tissue phenotypic features from each region are used to predict and localize matched biomarker expression measured from the same spatial region. This one-to-one ROI-level pairing preserves regional molecular specificity, in contrast to bulk-tissue based gene expression that aggregates signals across heterogeneous tissue compartments. By leveraging ROI-level spatial proteogenomic data across multiple biomarkers, it supports spatial modeling of broader biomarker expression programs rather than focused gene or protein panels. X-SPATIO integrates ROI-level biomarker expression from GeoMx DSP with weakly supervised MIL on matched H&E morphology, enabling interpretable discovery of which regions within each ROI encode high versus low expression of specific biomarkers. The pipeline combines segmentation-guided patch extraction, a custom attention-based MIL architecture, and spatial heatmap reconstruction to highlight the regions that are most informative for expression prediction and to visualize the morphological patterns that drive the model’s decisions.

By directly linking deep-learning and spatial proteogenomic profiling, X-SPATIO provides a practical software framework for both pathologists and computational researchers to interrogate how localized histologic patterns relate to underlying molecular states. This approach addresses a key limitation of existing H&E-based models which is the absence of spatially resolved molecular ground truth while retaining the scalability and accessibility of routine histopathology. As such, X-SPATIO provides a software framework for spatially informed biomarker analysis and model validation, by connecting routine histopathology with spatial molecular ground truth.

## MATERIALS AND METHODS

An overview of the X-SPATIO workflow is shown in Figure 1. In the following sections, we detail each stage of the pipeline, including ROI-aware segmentation, encoder-based feature extraction, spatially constrained multiple-instance learning, model evaluation, and attention-based spatial visualization.

**Figure 1.**
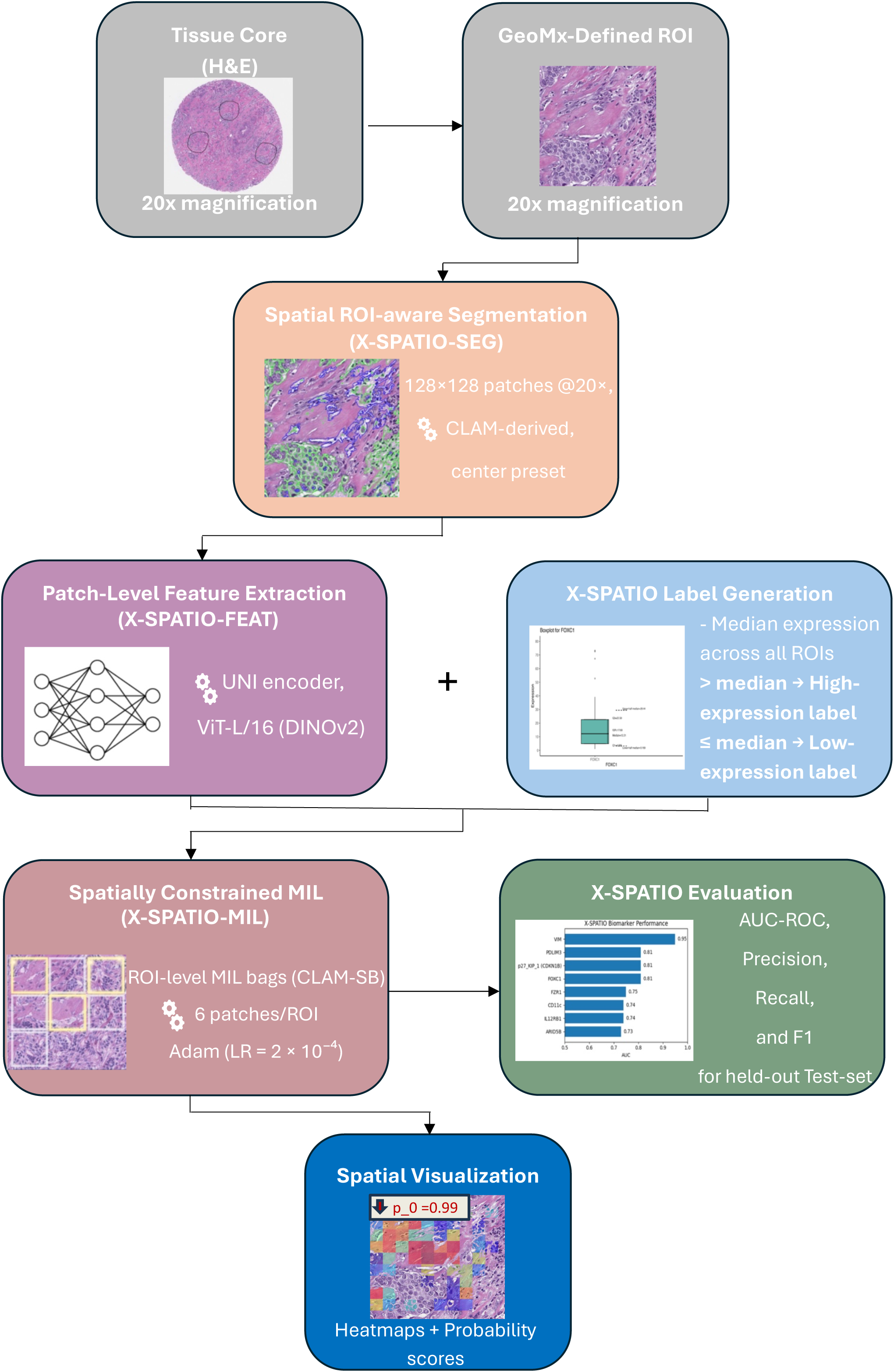
Overview of the X-SPATIO workflow.

### Dataset

GeoMx DSP dataset comprising two formalin-fixed paraffin-embedded (FFPE) TNBC tissue microarrays (TMAs: TMA1 and TMA2) with a total of 97 cores (1.5 mm cores, sectioned at 5 μm thickness) was analyzed, from which 272 pathologist-annotated ROIs were selected based on combined fluorescence (SYTO13 for DNA, PanCK, and CD45) and H&E morphology (Figures 2 and 3) and profiled using the Whole Transcriptome and Immuno-Oncology Proteome Atlases ^18,19^. RNA counts were normalized using Q3 method, while protein measurements were normalized to housekeeping targets, with normalization performed independently for each TMA to account for batch-specific effects (SpatioScope abstract submitted to ASIP’s Pathobiology 2026).

**Figure 2.**
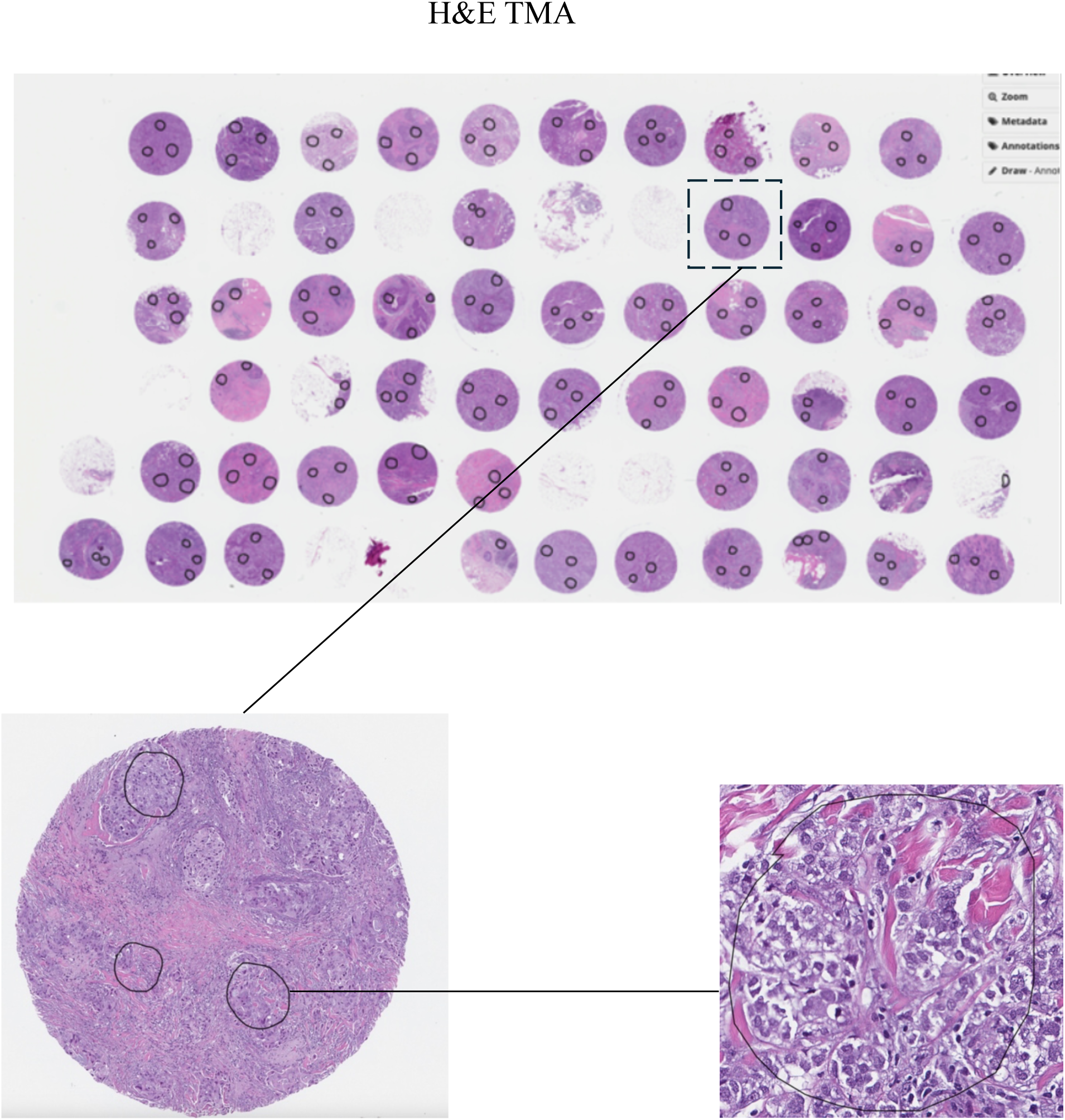
Hematoxylin and eosin (H&E) image of the tissue microarray (TMA 1) containing 57 invasive triple-negative breast cancer (TNBC) cores.

**Figure 3.**
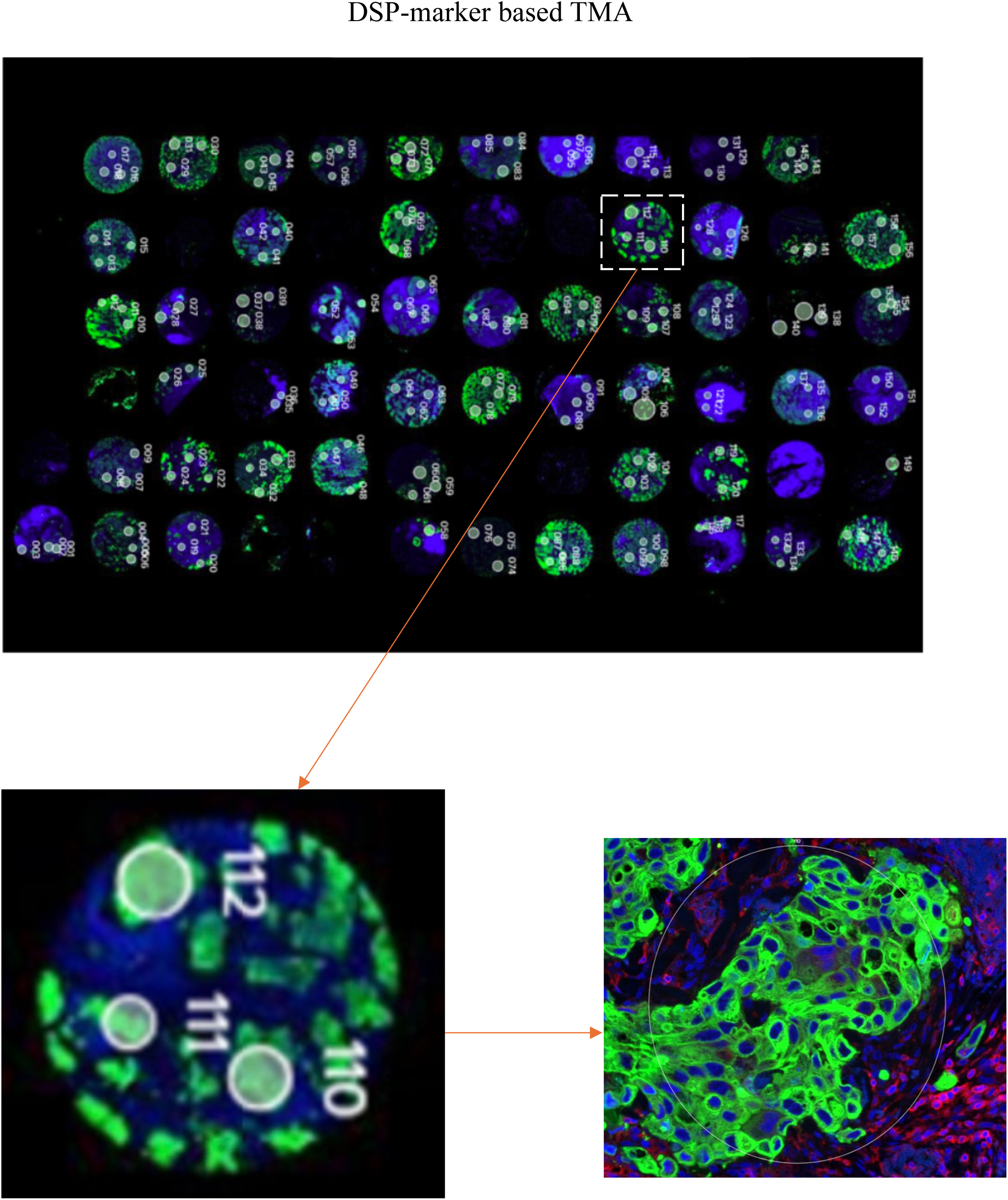
DSP marker-based visualization of the tissue microarray (TMA 1) containing 57 invasive triple-negative breast cancer (TNBC) cores.

To quantify overall variability of biomarker expression and determine their thresholds used in MIL training, global expression distributions were summarized using marker-wise boxplots depicting median, Q1, Q3, interquartile range, and half-median statistics (Figures 4A–4F).

**Figure 4.**
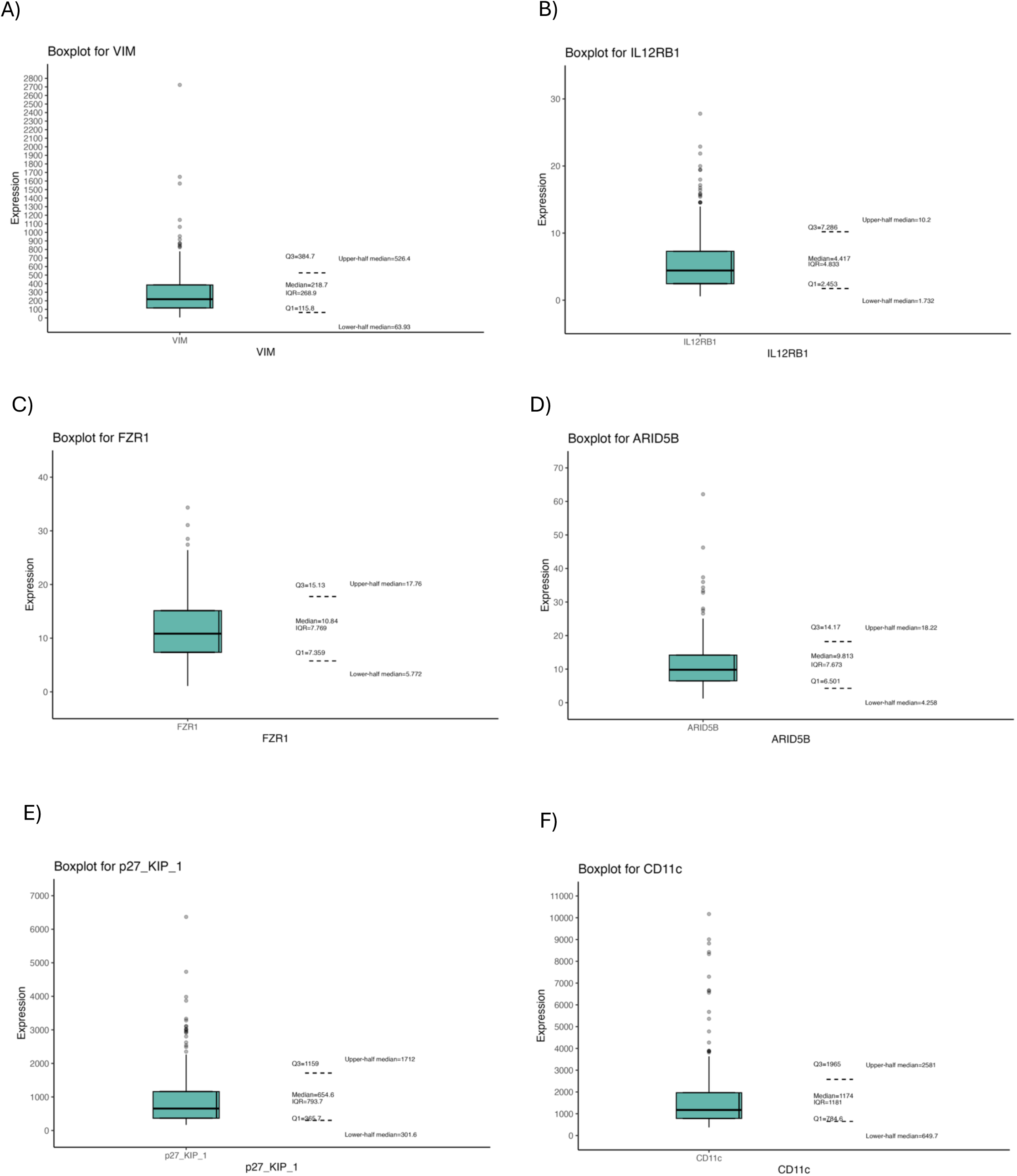
Boxplots summarizing expression levels for eight markers: (A) VIM, (B) IL12RB1, (C) FZR1, (D) ARID5B, (E) CDKN1B (p27_KIP_1) and (F) CD11c computed across all DSP ROIs independent of TMA core of origin. For each marker, boxplots display the median, first quartile (Q1), third quartile (Q3), interquartile range (IQR), and lower/upper half medians.

### Spatial ROI-Aware Segmentation (X-SPATIO-SEG)

Segmentation was performed using a CLAM-derived module adapted specifically for GeoMx ROI geometry ^11^. A center-weighted mask was applied to restrict tissue extraction to the defined ROI and to minimize inclusion of peripheral tissue, adjacent TMA cores, or folded areas. After masking, each ROI was tiled into non-overlapping 128 × 128-pixel patches at 20X magnification. Patches were retained only if they showed substantial spatial overlap with the ROI mask, ensuring that all downstream analyses were based solely on tissue regions that contributed to the spatial transcriptomic and proteomic measurements.

### Patch-Level Feature Extraction (X-SPATIO-FEAT)

Each retained ROI patch was converted into a high-dimensional feature embedding using the UNI histopathology encoder, a DINOv2-trained ViT-L/16 model ^20–22^. Patches (128 × 128-pixels at 20X) were independently encoded to capture local histopathologic features, including nuclear morphology, stromal structure, and epithelial organization. The feature extraction module is encoder-agnostic and supports substitution of alternative histology foundation models as new models emerge.

### Spatially Constrained MIL Modeling (X-SPATIO-MIL)

Each ROI was modeled as a multiple-instance learning bag of its constituent patches. Biomarker expression was binarized into high- and low-expression groups using median thresholds across all ROIs (Figure 4). X-SPATIO-MIL extends the CLAM-SB architecture by applying an attention-based MIL model restricted to ROI-specific patches to generate biomarker predictions and spatial attention maps^11^. The model was trained using a learning rate of 2 × 10⁻⁴ for 200 epochs with the Adam optimizer, a dropout rate of 0.25, and a hold-out cross-validation performed at 10 folds. During each training step, the CLAM-SB sampler was configured to select at least three patches per ROI as a set of candidate “positive or negative” instances used for learning.

### Model Evaluation

Model performance was assessed on held-out test (10%) ROIs using area under the receiver operating characteristic curve (AUC-ROC), accuracy, precision, recall, and F1-score.

### Spatial Visualization and Heatmap Generation

Attention weights were mapped back to patch coordinates within each ROI and rendered as continuous heatmaps overlaid on the respective H&E images. These heatmaps highlight tissue regions most strongly contributing to biomarker expression predictions and serve as the primary interpretability output of X-SPATIO.

## RESULTS

### Model Performance and AUC Summary

X-SPATIO demonstrated strong predictive performance across several biomarkers profiled by GeoMx DSP, with AUC-ROC values ranging from 0.79-0.97. Table 1 and Figure 5 summarize the performance metrics for each marker.

**Figure 5.**
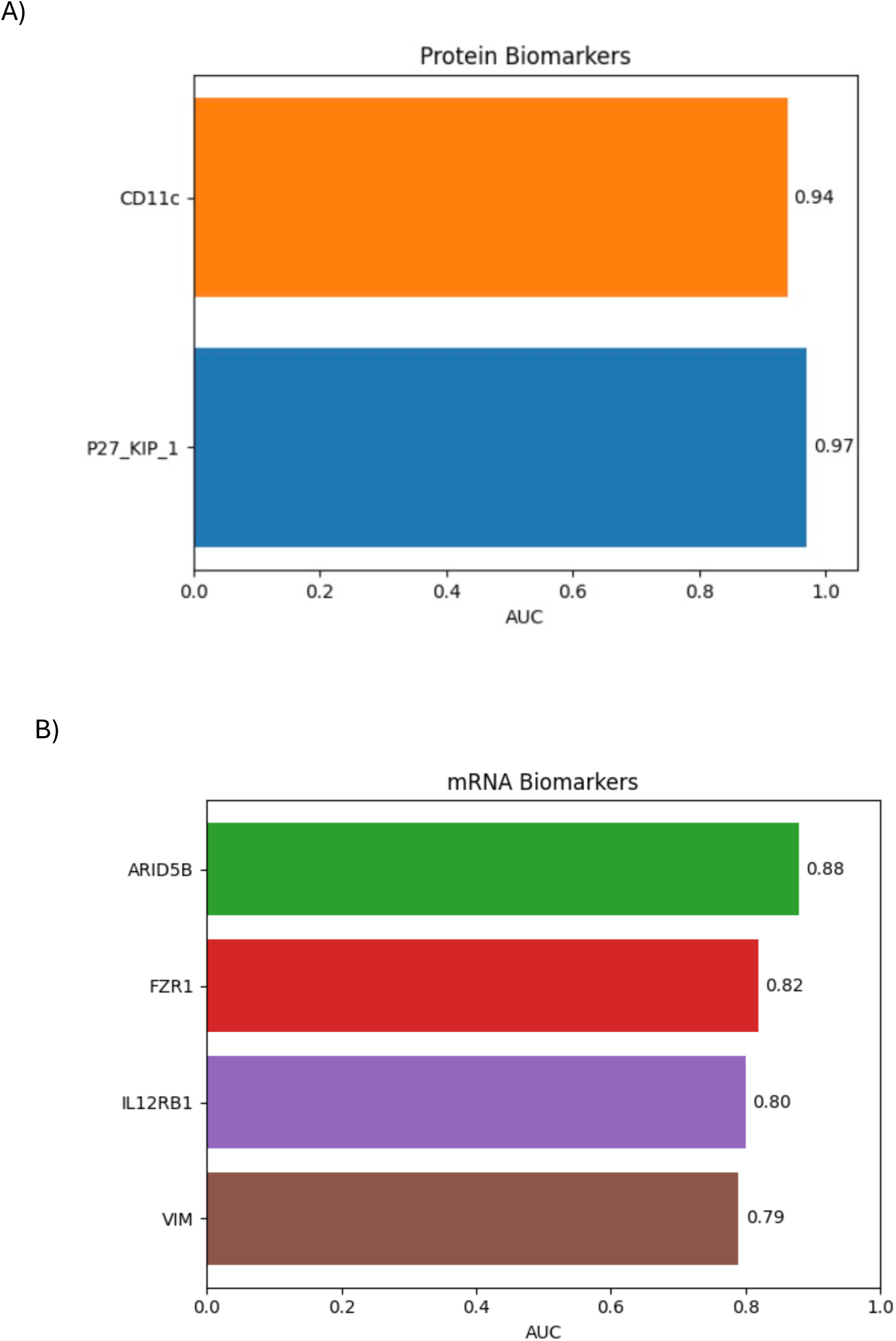
Bar plots showing X-SPATIO performance (AUC) for (A) protein (B) and mRNA biomarkers.

**Table 1:** Performance metrics for X-SPATIO biomarker prediction on held-out test sets, including area under the receiver operating characteristic curve (AUC), precision (positive predictive value), recall (sensitivity), F1 score (harmonic mean of precision and recall), with corresponding 95% confidence interval ranges (CI). Test Error represents the misclassification rate on the held-out test set.

| Biomarker | Expression<br>Source | AUC | AUC CI<br>range | Test<br>Error | Precision | Precision<br>CI Range | Recall | Recall<br>CI<br>Range | F1 | F1 CI<br>Range |
| --- | --- | --- | --- | --- | --- | --- | --- | --- | --- | --- |
| P27_KIP_1 | Protein | 0.97 | (0.89, 1) | 0.07 | 0.94 | (0.87, 1) | 0.91 | (0.82, 1) | 0.91 | (0.82, 1) |
| CD11c | Protein | 0.94 | (0.95, 1) | 0.07 | 0.94 | (0.87, 1) | 0.91 | (0.82, 1) | 0.91 | (0.82, 1) |
| ARID5B | mRNA | 0.88 | (0.64, 0.97) | 0.19 | 0.82 | (0.67, 0.97) | 0.81 | (0.67, 0.96) | 0.81 | (0.66, 0.96) |
| FZR1 | mRNA | 0.82 | (0.62, 0.96) | 0.19 | 0.83 | (0.68, 0.97) | 0.82 | (0.68, 0.96) | 0.82 | (0.68, 0.96) |
| IL12RB1 | mRNA | 0.8 | (0.76, 1) | 0.18 | 0.82 | (0.68, 0.97) | 0.82 | (0.68, 0.96) | 0.82 | (0.68, 0.96) |
| VIM | mRNA | 0.79 | (0.65, 0.98) | 0.18 | 0.82 | (0.68, 0.97) | 0.82 | (0.68, 0.96) | 0.82 | (0.67, 0.96) |

X-SPATIO generated biologically meaningful and spatially coherent attention maps across all evaluated TNBC biomarkers, accurately distinguishing high- and low-expression ROIs based solely on H&E morphology. Depending on the model’s association of the biomarkers’ expression with the tissue-morphological patterns, a biomarker could be predicted as either high or low-expressed in several ROI-patches, where red, orange and yellow-patches indicate regions that influence the model’s prediction the most, and green, cyan and blue-patches indicate regions that influence the prediction the least (Figure 6-11). The p_0 and p_1 indicate probabilities for the predicted class (high or low-expression) using top-10-scored patches and non-predicted class (low or high expression) using bottom-10-scored patches, respectively (Table 2). The following subsections illustrate representative examples for individual biomarkers, linking predicted expression with the corresponding histomorphologic patterns.

**Figure 6.**
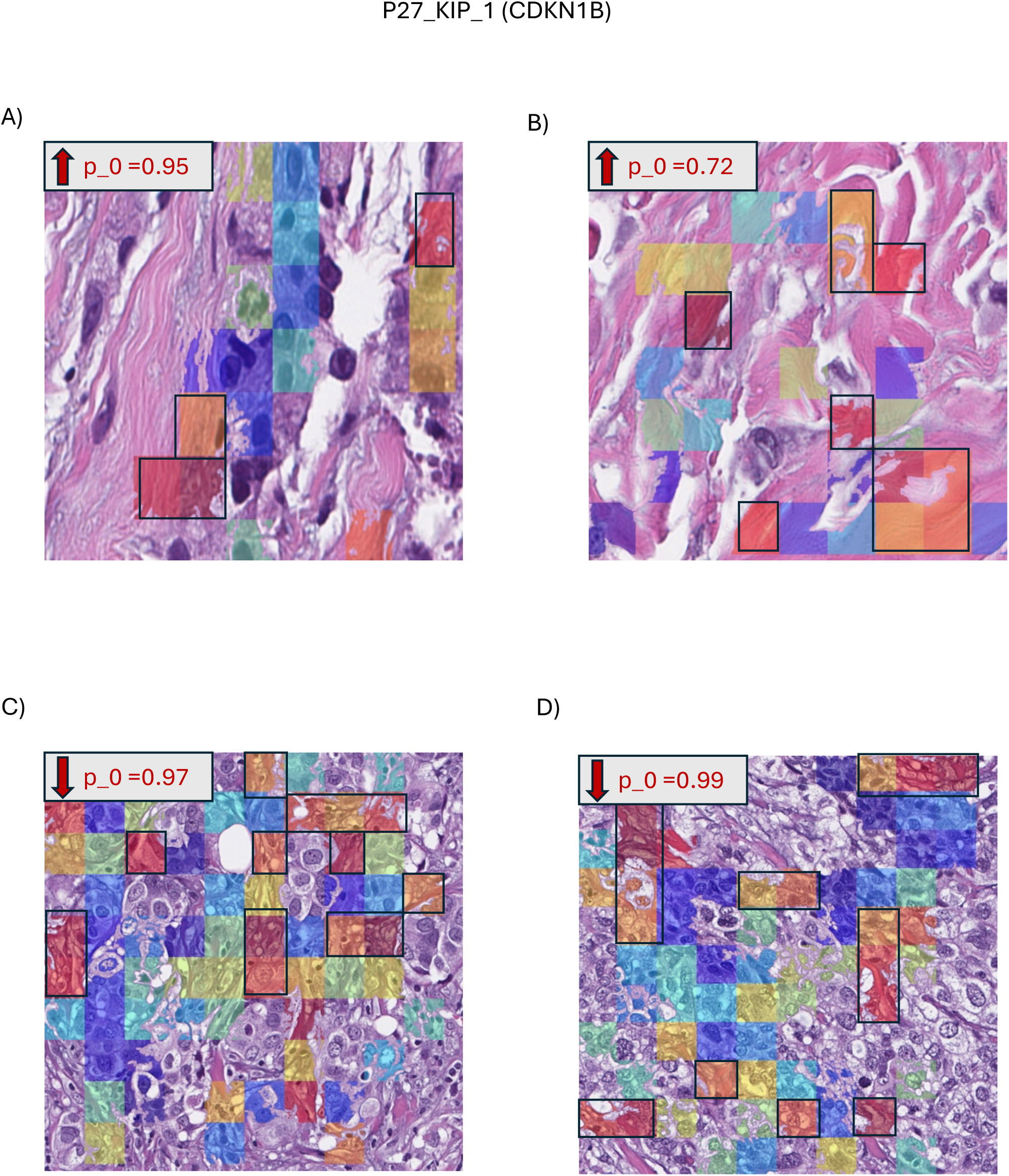
X-SPATIO attention heatmaps for CDKN1B high-expression ROIs (A) 134_TMA2, (B) 62_TMA2, and low-expression ROIs (C) 7_TMA1, (D) 49_TMA1. Red, orange, and yellow patches indicate regions that most strongly influence the model’s prediction, while green, cyan, and blue patches indicate regions with the least influence.

**Table 2:** X-SPATIO-predicted expression probabilities for biomarkers across TNBC ROIs for the held-out test set. For each ROI, the model predicts whether the biomarker is in high or low expression state; where pred_0 = Predicted expression class (high/low expression) using the top-10-scored attention patches; p_0 = Probability of the predicted class using top-10-scored attention patches; pred_1 = Predicted expression class (high/low) using the bottom-10-scored attention patches; p_1 = Probability of the predicted class using bottom-10-scored attention patches.

| Biomarker | Core | ROI_ID | pred_0 | p_0 | pred_1 | p_1 | Figure No |
| --- | --- | --- | --- | --- | --- | --- | --- |
| P27_KIP_1 | C46_TMA2 | 134_TMA2 | high_expression_genes | 0.95 | low_expression_genes | 0.05 | 6A |
|  | C25_TMA2 | 62_TMA2 | high_expression_genes | 0.72 | low_expression_genes | 0.28 | 6B |
|  | C4_TMA1 | 7_TMA1 | low_expression_genes | 0.97 | high_expression_genes | 0.03 | 6C |
|  | C23_TMA1 | 49_TMA1 | low_expression_genes | 0.99 | high_expression_genes | 0.01 | 6D |
| CD11c | C50_TMA2 | 146_TMA2 | high_expression_genes | 0.99 | low_expression_genes | 0.01 | 7A |
|  | C49_TMA2 | 142_TMA2 | high_expression_genes | 0.99 | low_expression_genes | 0.01 | 7B |
|  | C10_TMA1 | 23_TMA1 | low_expression_genes | 0.99 | high_expression_genes | 0.01 | 7C |
|  | C47_TMA1 | 106_TMA1 | low_expression_genes | 0.98 | high_expression_genes | 0.02 | 7D |
| ARID5B | C35_TMA2 | 91_TMA2 | high_expression_genes | 0.6 | low_expression_genes | 0.4 | 8A |
|  | C49_TMA2 | 142_TMA2 | high_expression_genes | 0.73 | low_expression_genes | 0.27 | 8B |
|  | C47_TMA2 | 138_TMA2 | low_expression_genes | 0.92 | high_expression_genes | 0.08 | 8C |
|  | C10_TMA1 | 23_TMA1 | low_expression_genes | 0.95 | high_expression_genes | 0.05 | 8D |
| FZR1 | C25_TMA2 | 61_TMA2 | high_expression_genes | 0.78 | low_expression_genes | 0.22 | 9A |
|  | C14_TMA2 | 34_TMA2 | high_expression_genes | 0.71 | low_expression_genes | 0.29 | 9B |
|  | C30_TMA1 | 66_TMA1 | low_expression_genes | 0.97 | high_expression_genes | 0.03 | 9C |
|  | C24_TMA1 | 53_TMA1 | low_expression_genes | 0.96 | high_expression_genes | 0.04 | 9D |
| IL12RB1 | C50_TMA2 | 147_TMA2 | high_expression_genes | 0.66 | low_expression_genes | 0.34 | 10A |
|  | C47_TMA1 | 106_TMA1 | high_expression_genes | 0.53 | low_expression_genes | 0.47 | 10B |
|  | C10_TMA1 | 23_TMA1 | low_expression_genes | 0.63 | high_expression_genes | 0.37 | 10C |
|  | C26_TMA1 | 56_TMA1 | low_expression_genes | 0.66 | high_expression_genes | 0.34 | 10D |
| VIM | C50_TMA2 | 147_TMA2 | high_expression_genes | 0.75 | low_expression_genes | 0.25 | 11A |
|  | C53_TMA2 | 159_TMA2 | high_expression_genes | 0.84 | low_expression_genes | 0.16 | 11B |
|  | C24_TMA1 | 53_TMA1 | low_expression_genes | 0.8 | high_expression_genes | 0.2 | 11C |
|  | C16_TMA1 | 33_TMA1 | low_expression_genes | 0.6 | high_expression_genes | 0.4 | 11D |

Across a diverse set of tumor-intrinsic, stroma, immune, and cell-cycle–associated biomarkers, X-SPATIO produced spatial attention patterns that were biologically coherent and consistent with known compartment-specific morphologies in TNBC. p27KIP1 (CDKN1B), a G1 cell-cycle inhibitor, exhibited attention in high-expression tumors over fibrous stroma and low-cellularity epithelial regions lacking nuclear crowding or mitotic figures, consistent with preserved cell-cycle inhibition (Figure 6A, 6B), while CDKN1B-low tumors showed attention localized to densely cellular tumor regions with crowded, irregular nuclei and architectural disorganization, reflecting increased proliferative activity (Figure 6C, 6D) ^23,24^.

CD11c (ITGAX), a marker of antigen-presenting myeloid cells, showed attention in CD11c-high tumors over cellular stromal regions enriched with larger immune cells and stroma–tumor interfaces, consistent with a myeloid-inflamed microenvironment (Figure 7A, 7B), whereas CD11c-low tumors demonstrated attention over dense, collagen-rich fibrous stroma with minimal stromal cellularity, indicative of a myeloid-poor, immune-cold state (Figure 7C, 7D) ^25,26^.

**Figure 7.**
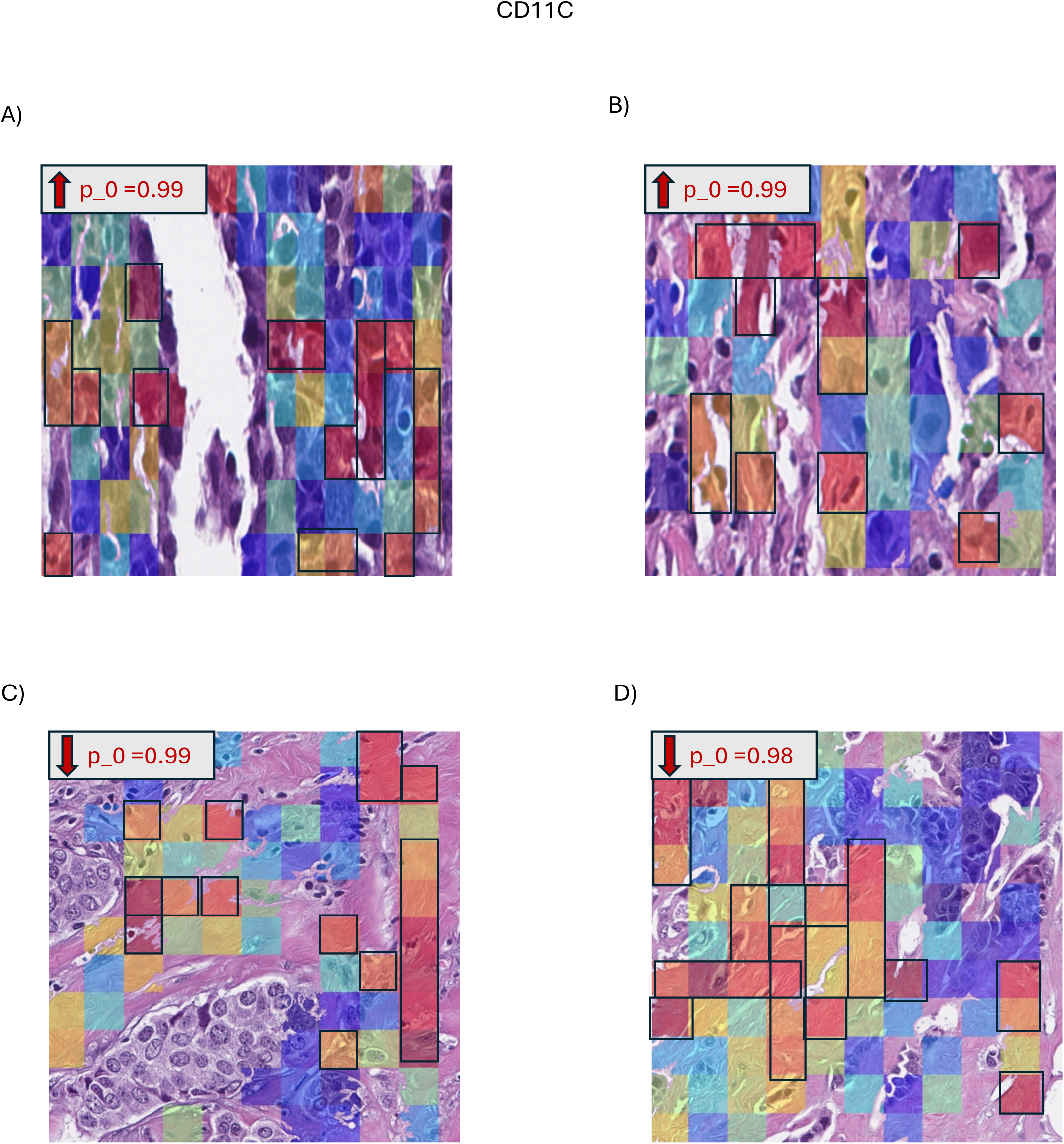
X-SPATIO attention heatmaps for CD11C high-expression ROIs (A) 146_TMA2, (B) 142_TMA2 and low-expression ROIs (C) 23_TMA1 (D) 106_TMA1. Red, orange, and yellow patches indicate regions that most strongly influence the model’s prediction, while green, cyan, and blue patches indicate regions with the least influence.

ARID5B, a chromatin-associated transcriptional regulator implicated in stromal and immune-related programs, demonstrated attention in high-expression tumors localized primarily to dense fibrous stromal regions and stroma–tumor interfaces (Figure 8A, 8B), while ARID5B-low tumors exhibited attention over uniform, collagen-rich fibrous stroma with minimal cellular or interface complexity, consistent with reduced stroma or immune-associated transcriptional activity (Figure 8C, 8D) ^27-29^.

**Figure 8.**
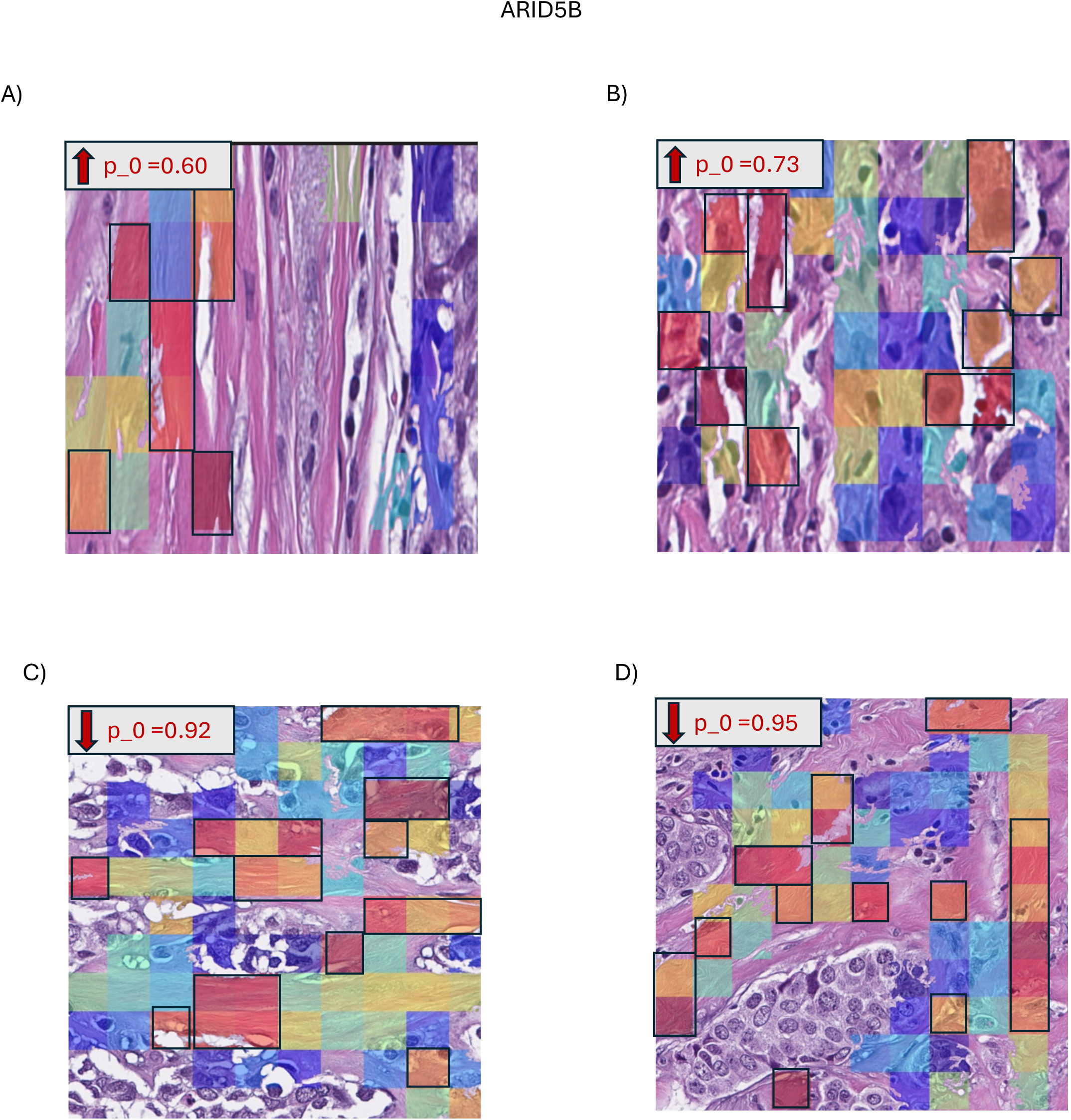
X-SPATIO attention heatmaps for ARID5B high-expression ROIs (A) 91_TMA2, (B) 142_TMA2 and low-expression ROIs (C) 138_TMA2, (D) 23_TMA1. Red, orange, and yellow patches indicate regions that most strongly influence the model’s prediction, while green, cyan, and blue patches indicate regions with the least influence.

FZR1 (Cdh1), a key regulator of Anaphase-Promoting Complex/Cyclosome (APC/C)-mediated cell-cycle control, exhibited attention in FZR1-high tumors over fibrous stroma and epithelial tumor cells with uniform, round nuclei, consistent with restrained proliferation (Figure 9A, 9B), whereas FZR1-low tumors showed attention concentrated in densely packed, pleomorphic tumor regions with nuclear crowding and architectural irregularity, reflecting loss of cell-cycle regulation (Figure 9C, 9D) ^30,31^.

**Figure 9.**
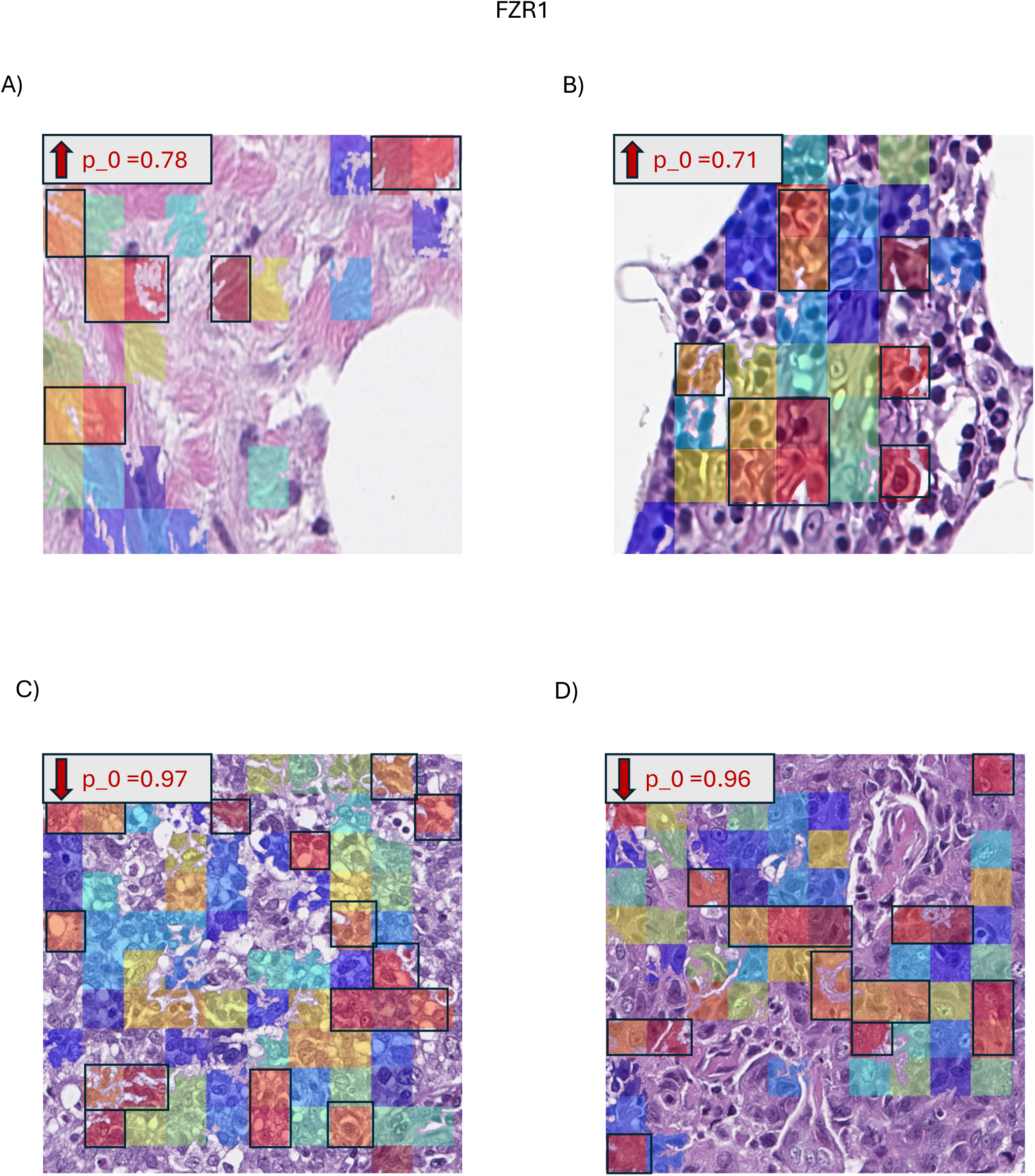
X-SPATIO attention heatmaps for FZR1 high-expression ROIs (A) 61_TMA2, (B) 34_TMA2, and low-expression ROIs (C) 66_TMA1, (D) 53_TMA1. Red, orange, and yellow patches indicate regions that most strongly influence the model’s prediction, while green, cyan, and blue patches indicate regions with the least influence.

IL12RB1, an immune-associated receptor expressed primarily by activated immune cells, demonstrated attention in high-expression tumors over cellular stromal regions containing scattered immune cells embedded within fibrous matrix, consistent with an immune-active microenvironment (Figure 10A, 10B), while IL12RB1-low tumors showed attention localized to dense, hypocellular fibrous stroma with minimal immune infiltration, indicative of an immune-cold state (Figure 10C, 10D) ^32,33^.

**Figure 10.**
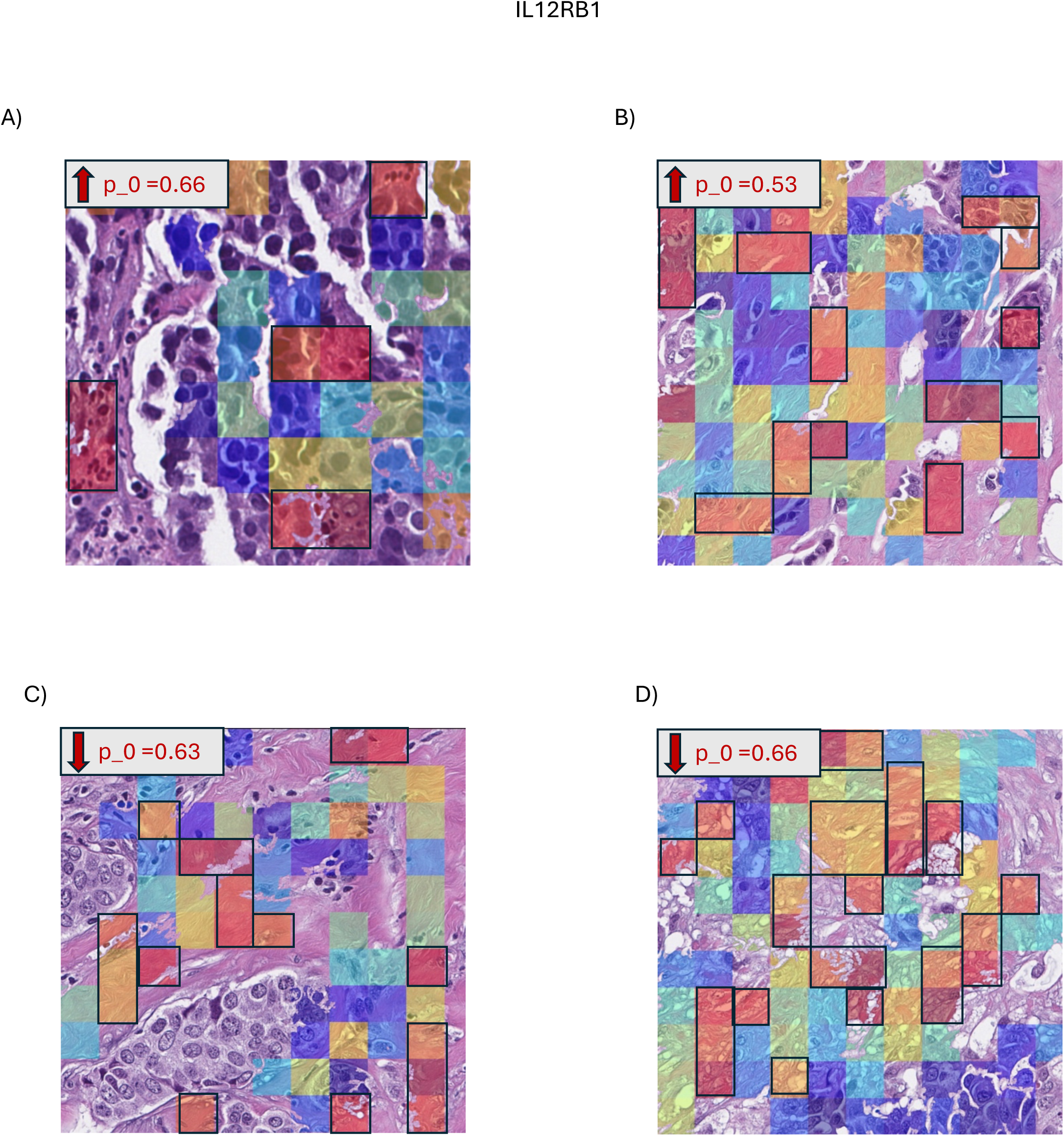
X-SPATIO attention heatmaps for IL12RB1 high-expression ROIs (A) 147_TMA2, (B) 106_TMA1 and low-expression ROIs (C) 23_TMA1, (D) 56_TMA1. Red, orange, and yellow patches indicate regions that most strongly influence the model’s prediction, while green, cyan, and blue patches indicate regions with the least influence.

Vimentin (VIM), a canonical marker of EMT in TNBC, showed attention in VIM-high tumors localized to poorly cohesive tumor cells with elongated morphology, nuclear crowding and architectural breakdown, consistent with mesenchymal transformation (Figure 11A, 11B), whereas VIM-low tumors exhibited attention over cohesive epithelial tumor regions or adjacent fibrous stroma, reflecting preserved epithelial architecture and reduced EMT features (Figure 11C, 11D) ^34-36^. Collectively, these results demonstrate that X-SPATIO captures spatially localized, pathway-relevant morphologic cues across multiple biological axes.

**Figure 11.**
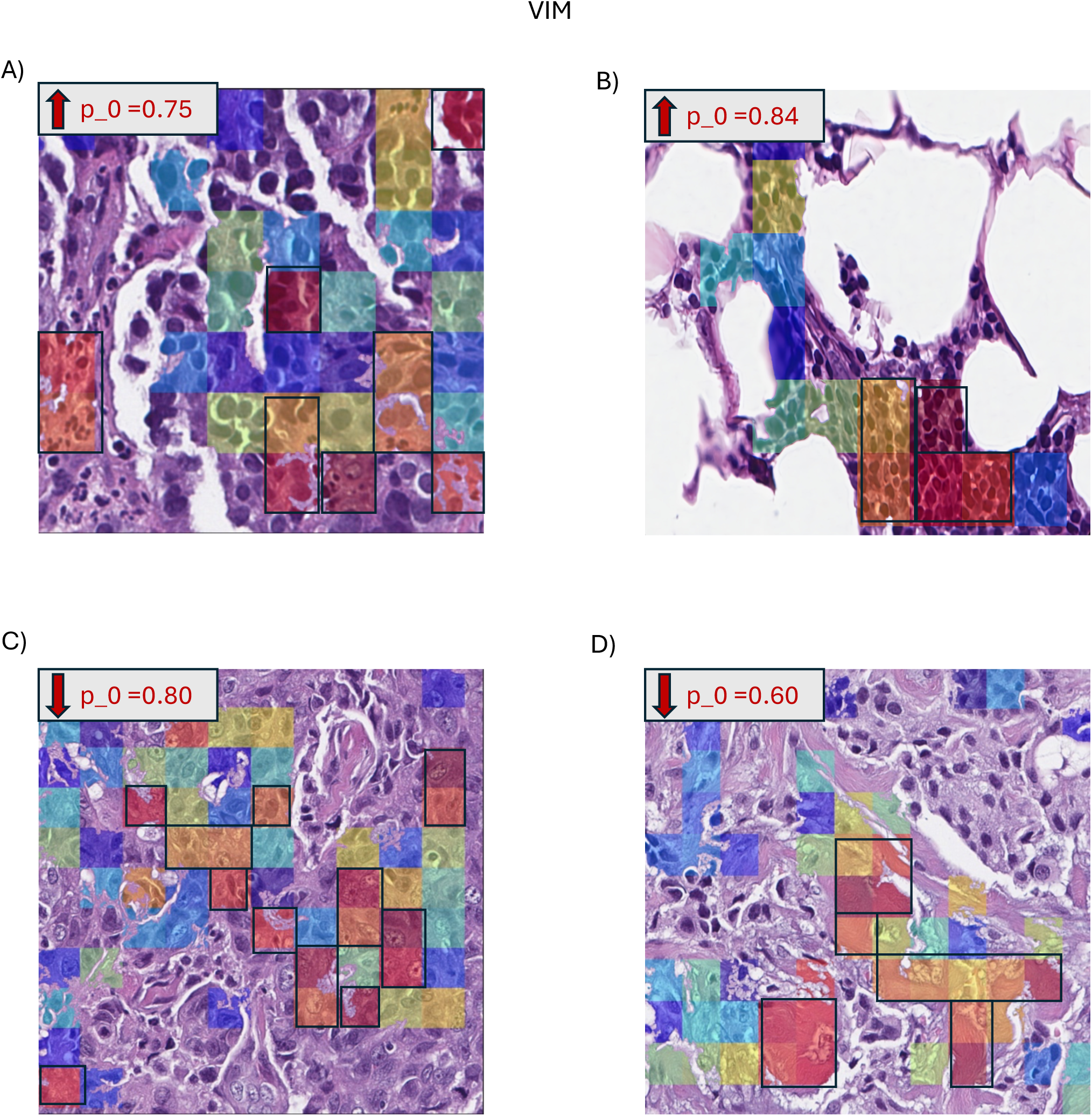
X-SPATIO attention heatmaps for VIM high-expression ROIs (A) 147_TMA2, (B) 159_TMA2, and low-expression ROIs (C) 53_TMA1, (D) 33_TMA1. Red, orange, and yellow patches indicate regions that most strongly influence the model’s prediction, while green, cyan, and blue patches indicate regions with the least influence.

## DISCUSSION

In this study, we introduce X-SPATIO, an explanatory deep learning pipeline that links H&E-defined tumor ROIs with region-matched transcriptomic and proteomic data. By leveraging ROI-specific expression across multiple biomarkers as ground truth, X-SPATIO captures morpho-molecular associations that are otherwise lost by bulk tissue-based gene expression. X-SPATIO’s attention maps reveal tumor regions that are predictive of spatially distinct biomarker expression in TNBC. X-SPATIO achieved strong predictive performance (AUC = [0.79, 0.97], 95% CI range = [0.62, 1.0]) and enabled interpretations of morpho-molecular attention maps that revealed distinct and biologically coherent spatial patterns across multiple biomarkers, reflecting both tumor-intrinsic and microenvironmental molecular programs.

High expression of cell-cycle regulators CDKN1B (p27KIP1) and FZR1 localized attention to fibrous stroma and epithelial regions with uniform nuclear morphology and reduced cellular density, consistent with preserved cell-cycle control, whereas low expression concentrated attention in densely packed tumor regions with nuclear crowding and pleomorphism reflecting increased proliferation. Immune-associated markers CD11c and IL12RB1 showed similar expression-dependent spatial patterns, with high expression highlighting immune-enriched stromal compartments and stroma–tumor interfaces indicative of immune-inflamed microenvironments, and low expression localizing to hypocellular, collagen-rich fibrous stroma characteristic of immune-cold states. ARID5B exhibited analogous behavior, with high expression emphasizing dense stromal regions and interface zones suggestive of active stromal–immune signaling, and low expression associated with uniform fibrotic architecture and minimal cellular complexity. Finally, VIM expression patterns distinguished mesenchymal from epithelial phenotypes, with VIM-high tumors showing attention over poorly cohesive, elongated tumor cells and architectural breakdown, while VIM-low tumors retained cohesive epithelial morphology. Collectively, these expression-linked spatial patterns demonstrate that X-SPATIO captures pathway-relevant morphologic cues across multiple biological axes, providing spatially grounded, biologically interpretable insights into how molecular programs manifest within TNBC tumor and microenvironmental architecture.

Most deep learning models trained on whole-slide labels lack the ability to assess predictions at the sub-region level, limiting insight into whether learned features correspond to true spatial biology. X-SPATIO integrates ground-truth spatial expression to supervise model attention at the ROI level, enabling direct validation of prediction localization within tissue ROIs. This sub-regional spatial supervision distinguishes X-SPATIO from conventional slide-level approaches by explicitly linking histomorphology to region-matched biomarker expressions. As a result, model outputs become spatial hypotheses that can be interrogated experimentally, increasing confidence in computational approaches for biomarker discovery and ROI-level interpretation.

The X-SPATIO framework further enables identification of biomarkers whose spatial expression is strongly encoded in histomorphology, as well as those exhibiting weak or inconsistent attention patterns across tumor regions. Biomarkers with borderline predictive performance likely correspond to molecular programs that are either not readily observable in H&E-morphology or require larger and more diverse datasets for stable learning. Recognizing these limitations helps prevent over-interpretation and underscores the complementary roles of computational modeling and experimental spatial profiling in characterizing tumor heterogeneity.

Future work will extend X-SPATIO to a substantially larger TNBC cohort, increasing both morphologic and molecular diversity and enabling more robust assessment of biomarker-specific performance and generalizability. As these expanded datasets become available, the framework is expected to support more stable learning of subtle morpho–molecular associations and improved spatial consistency of model predictions. In addition, the integration of single-cell profiling data may further enhance model robustness by refining cell-type–specific signals and providing higher-resolution molecular context for spatial interpretation.

In summary, X-SPATIO serves as a first-of-its-kind explanatory software framework that links histopathology with spatially measured biomarker expression, enabling cost-effective and scalable inference across a broad range of genes and proteins while providing sub-regional spatial interpretability not previously achievable in multiple-instance learning approaches. X-SPATIO offers a streamlined, reproducible workflow that supports rapid deployment and is readily generalizable across cancer types. Together, these capabilities position X-SPATIO as a foundational Artificial Intelligence (AI) tool for biologically grounded biomarker discovery, particularly in clinically challenging cancer types.

## Data Availability Statement

All authors confirm that all data and code used in this study are publicly available at the GitHub repository: https://github.com/skr1/Xspatio (accessed January 31^st^, 2025).

## Conflict of Interest

No conflict of interest is reported by any of the authors.

## Funding

N/A

## Acknowledgements

The authors acknowledge the support of the Pathology Shared Resource (RRID: SCR_023479) at the Dartmouth Cancer Center (DCC; NCI Cancer Center Support Grant 5P30 CA023108-37) and the Friends of DCC for the generation of Digital Spatial Profiler Dataset.

## Author Contributions Statement

Conceptualization SSS; Data curation AAW, SSS, CCB, MDC, GJZ; Data generation SP, MKS; Formal analysis VRR, SSS, MKS (GeoMx dataset); Funding acquisition SSS (self for software development, implementation and runs), and SSS, CCB, MDC, GJZ, TAM and LJV (dataset); Investigation Methodology SSS, CCB, GJZ, VRR; Project administration SSS, GJZ; Resources SSS, MKS, SMP, GJZ, LJV; Software SSS, VRR; Supervision SSS (software development and implementation), GJZ (GeoMx); Validation SSS, VRR, CCB, GJZ and MDC; Visualization SSS, VRR; Writing – original draft VRR, SSS; Writing – review & editing SSS, VRR, GJZ, MKS, CCB, AAW, MDC, SMP, TAM, XL and LJV

## Notes

### Competing Interest Statement

The authors have declared no competing interest.

